# Semi-Automated Identification of Ontological Labels in the Biomedical Literature with goldi

**DOI:** 10.1101/073460

**Authors:** Christopher B. Cole, Sejal Patel, Leon French, Jo Knight

## Abstract

Recent growth in both the scale and the scope of large publicly available ontologies has spurred the development of computational methodologies which can leverage structured information to answer important questions. However, ontological labels, or “terms” have thus far proved difficult to use in practice; text mining, one crucial aspect of electronically understanding and parsing the biomedical literature, has historically had difficulty identifying “terms” in literature. In this article, we present goldi, an open source R package whose goal it is to identify terms of variable length in free form text. It is available at https://github.com/Chris1221/goldi or through CRAN. The algorithm works through identifying words or synonyms of words present in individual terms and comparing the number of present words to an acceptance function for decision making. In this article we present the theoretical rationale behind the algorithm, as well as practical advice for its usage applied to Gene Ontology term identification and quantification. We additionally detail the options available and describe their respective computational efficiencies.

## Introduction

Large Ontologies (LO) classify information and entities into meaningful and occasionally hierarchical categories. [1] The use of these ontologies has been a hot topic in recent research; these databases may be leveraged to answer pressing questions in various fields. Gene Ontology (GO), a platform which allows researchers to describe gene function, among other things, through a structured and hierarchical vocabulary, is one such ontology which has seen much recent interest. [2] The terms have a complex relational structure, and their specificity generally scales with their length. There has been recent interest in identifying these “terms” or gene functions in the biomedical literature for use in comparative analyses and machine annotation. [3–5] However, the automated discovery and subsequent usage of ontological labels, or “terms” in literature mining, specifically in the biomedical literature, has historically been difficult. [6] Issues in term identification range from field specific jargon to inefficient computational algorithms used to parse data. The presence of this rich and detailed information presents researchers with the opportunity to delve deeply into biological relationships, yet historically Information Extraction (IE) has been a stumbling block in research pipelines.

To understand and apply the information present in GO, a researcher may need to identify the terms in literature. For example, researchers may wish to find terms which are enriched over chance in a particular field in a literature base when compared to a related field. However, given this problem, a researcher would run into several issues:

- Identify co-occurring words which constitute a “term”.
- Efficiently computing, summarizing, and reporting associations across a large corpus of literature.
- Implementing strict quality control in order to reduce the chances of false identifying terms or missing truly present terms.
- Efficiently deal with issues of polysemy (multiple meanings for the same word), synonymy (multiple synonyms or different words with the same meaning), and homosemy (multiple words having the same spelling but different meanings).

In this article we present goldi as an R package capable of addressing these concerns and performing multi-word term identification and quantification in natural language.

## Design and Implementation

A large issue in term recognition thus far, and a large part of our design philosophy in creating goldi, has been the relatively large divide between computational methodologies and expert knowledge which is needed to understand free text. Though this is a significant issue in all fields, it is especially apparent in biology and genetics where non-dictionary terms and acronyms form a large portion of the literature’s content; computational algorithms struggle without a clear cut set of rules in these cases. [4]

`goldi` attempts to address these concerns by utilizing human knowledge and domain specific information to function optimally. As an example, the strength and flexibility of goldi is greatly enhanced by user constructed lists of synonyms. Though this requires a larger time investment up front, the benefits are large. Furthermore, we provide an avenue for users to share their lists of synonyms on our Github project site wiki.

`goldi` works with a list of terms, or labels, which the user must supply. For this publication, we work with all GO molecular function terms. These terms will be identified and quantified in the corpus of literature, which represents the sample space for the problem. goldi will identify terms in the corpus and return their positions along with context. From this the user can easily compute various metrics of interest, including frequency in subsets and weighting importance by inverse frequency.

The computational algorithm proceeds according to the following:

- Read in text and term list to internal memory.
- Perform quality control on text and term list
- Construct term document matrices (TDM) for both text and term list.
- Integrate given synonyms into the term document matrices.
- Match terms from the term list to the text and compare to the acceptance function 𝒜(*l*) given term length.
- Return matched terms and their contexts in the article.

Going into slightly more detail, we describe each step in turn and explain its rationale.

Text from documents is read into the program in one of four ways. These options are summarized in **Table 1.** Either input is given as a character vector already read into R, or it is given as a file path. goldi performs a best guess approximation of what the user wants if the option is not specified. If the text is given as an already existing text object, this object is used for the rest of the analysis.

**Table 1.** Available options for reading in text to goldi.

| Argument | Description | Dependencies |
| --- | --- | --- |
| "local" | Uses a local character vector for the text document. No external reading is performed, and goldi assumes that the text is in the format which you prefer. | None |
| "txt" | Given a supplied file path, `base::readLines()` is used to read in the text and store in memory. | None |
| "R" | Given a supplied file path, `pdftools::pdf_txt()` is used to parse a PDF file and store the text data as a character vector in R. | pdftools |
| "py" | Given a supplied file path, `pdfminer` is used to parse a multi column PDF file and store the text data as a character vector in R. | pdftools |

If given as a file path, the text is read in one of three ways. Either the text is a text document (extension `.txt`) requiring no parsing, or it is a PDF file which must be scraped. In the former case, base R functions are used to read the text from the document. In the latter case, the user has two options to read the data. The first is `pdf_txt`() from the `pdftools` package, which is the faster option. However, this program is incapable of reading PDF files which have multiple columns. In this more complex case, the user may specify that the python program `pdfminer` is to be used. This program is included in the download of `goldi`.

Once the text has been read into internal memory, slight modifications are performed to prepare it for further analysis. First, vestiges from page breaks are removed and paragraphs are split into individual element sentences by terminal punctuation marks. Second, quality control is implemented on the text in order to standardize input and improve the accuracy of the program. These include the following commonly used steps:

- Case standardization
- Punctuation elimination (with exception cases)
- Numerical elimination (with exception cases)
- Removing of Stop Words (for a given language, or standard English by default)
- Consistent word stemming
- White space elimination and consideration

Quality control of text input is accomplished through the tm package. [7] This step is necessary to distinguish various forms of words and to standardize input.

Once the quality control has been performed on the text source, a term document matrix (TDM) is created. This matrix has columns for each sentence *i* and rows for each word *j* present in the whole document. Entries in the matrix are the number of times word *j* occurs in sentence *i*. A similar transformation is applied to both the term list and the any synonyms which are presented to `goldi`. This is in order to standardize the quality control procedures to facilitate comparisons.

`goldi` loops through each term in the term list, identifying how many of the words present in the term are also present in each sentence of the text document. If enough of the term’s words or synonyms are found in any of the sentences of the document, the term is declared to be present. The necessary number of words which must be present is a function of term length *n*, and given by a modifiable acceptance function 𝒜(*n*).

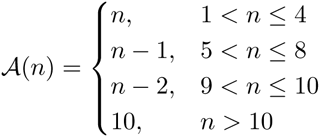

If the number of words from the term present in a particular sentence equal or exceed 𝒜(*n*), an association is declared.

The acceptance threshold can be fed to goldi as a vector of length *m* where *m* is the maximal term length that should be identified. The above acceptance function would be given in the vector `c(1,2,3,4,4,5,6,7,7,8,10)` for terms of up to 11 words in length.

Once these matches have been identified, the positions of the phrases used to identify them are recorded, and presented back to the user along with the number of times the term was identified in the article, if at all.

`goldi` follows 𝒪(2^*N*^) time, which may be mitigated by grouping terms into small chunks and processing in parallel.

Users may refer to both the supplemental material of this publication and the vignette for information on installation and execution of unit tests. We also present a simplified analysis in the vignettes which may be useful for beginners to natural language processing. `goldi` is written in `R` and `C++` integrated through `Rcpp` and `RcppArmadillo. goldi` imports several libraries including `tm, dplyr, SnowballC, foreach, mcSnow` and others. [8–15] In the supplemental material we give known stable versions of these packages.

goldi contains one overarching function `goldi::goldi()` which takes two mandatory arguments, `doc` and `terms`, containing, as mentioned above, either an R object or the path to a `.pdf` or `.txt` file and a newline delimited list of terms to be identified. A brief example is given below in Usage Instructions and a more complete run-through with examples and explanation is given in the “Basic Usage” vignette, also supplemental methods.

`goldi` is designed to be used in batch operations, however for smaller scale or pilot testing, an interaction version may be used, in which the program guides the user through the logic and process of designing their terms and synonyms. This may be performed by giving ‘‘`interactive`’’ to any of `lims, syn`, which will call `goldi::make.lims, goldi::make.syn`.

Optionally, a log file may be written to the path specified to the log `log = ‘‘path’’`, or if `log = TRUE` then `goldi` will write to `goldi.txt` in the current working directory. Logging is done through `futile.logger`. [9]

## Quality Assurance

goldi implements several layers of quality control. Unit testing is conducted on essential functions through the `testthat` package and test coverage estimated through covr. Additionally, continuous integration is tested for each new release by Travis CI on OSX and linux, and through Appveyor for windows.

## Results

We present the results of an example analysis, outlined in the supplied supplemental material and package vignettes, attempting to show terms which are over represented in a target group of Pubmed abstracts. For this section, we attempt to identify molecular function terms over represented in a query of 111 abstracts published between 2014 and 2015 identified with the search phrase “anaphylaxis genetics”. We use 1000 control abstracts identified through the phrase “immunology genetics” published between 2014 and 2015.

We use an analogue of the gene over expression analysis implemented by GOrilla to assess significant enrichment. Given *N* total abstracts and *B* of these annotated to a given GO term, the probability that *b* of the *n* abstracts in the target set are annotated to the same GO term is given by the tail of the Hypergeometric distribution:

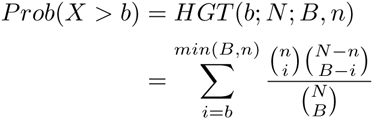

We fetch molecular function terms from the Gene Ontology website http://geneontology.org/ using a SQL query tool named GOOSE to obtain terms from the data base directly. Researchers could alternatively use the Quick GO website (http://www.ebi.ac.uk/QuickGO/) to extract GO terms for a set of target genes.

We fetch articles with the `RISmed` package in `R`.

After annotating each abstract with molecular function terms, we identify nine over expressed terms in the target set, as shown in **Table 2**. We provide a convenience function (`goldi::enrichment`) which recreates this analysis for the user given `goldi` input, though we also work through the code in the supplied vignettes.

**Table 2.** Terms enriched in abstracts matching “anaphylaxis genetics” Pubmed abstracts from 2014 to 2015 compared against “immunology genetics“ abstracts from the same period. Given *N* abstracts total and *B* identifications of the term, let *n* be the number of abstracts in the target set and *b* be the number of identifications in the target set. Following this, enrichment is defined as 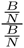while the P value is calculated through the hypergeometric tail *HGT* (*b*; *N*; *B*; *n*). P values adjusted through FDR correction to 5% false discovery rate. Only terms with more than five occurrences in the target set were analysed to reduce spurious associations.

| Term | Enrichment | P |
| --- | --- | --- |
| protein_C_(activated)_activity | 66.55 | $1.25 \times 10^{-20}$ |
| CD27_receptor_activity | 66.00 | $2.61 \times 10^{-21}$ |
| CD40_receptor_activity | 66.00 | $2.61 \times 10^{-21}$ |
| receptor_activator_activity | 66.00 | $2.61 \times 10^{-21}$ |
| receptor_activity | 66.00 | $2.61 \times 10^{-21}$ |
| IgE_binding | 65.34 | $8.63 \times 10^{-20}$ |
| kinase_activator_activity | 65.34 | $8.64 \times 10^{-20}$ |
| kinase_activity | 65.34 | $8.63 \times 10^{-20}$ |
| B_cell_receptor_activity | 62.22 | $5.11 \times 10^{-14}$ |
| T_cell_receptor_activity | 62.22 | $5.11 \times 10^{-14}$ |

The analysis took approximately three hours on a computer cluster, though all calculations were performed in serial.

## Availability and Future Directions

`goldi` is open source and released under an MIT license on Github at https://github.com/Chris1221/goldi. Additional documentation may be found in the wiki articles and vignettes of the package. goldi may be installed from CRAN with `install.packages("goldi")` or from Github with `devtools::install_github("Chris1221/goldi")`.

Typical and expected use of `goldi` is in the mining of biomedical literature, specifically through Gene Ontology terms. However, we see potential for this software outside of biology in fields such as the humanities and social media text mining. We believe that the identification of key terms is a fairly general process which may be applied in many areas, many of which we are unaware. Additionally, the code contained in this package may be included in different applications and software to incorporate the method into new areas.

## Conclusion

Here we present goldi, an R package to assist in the rapid identification and quantification of multi word terms in the literature. We have presented a use case scenario and hope that users will contribute their own research to our project page. We have released our software under an MIT license at https://github.com/Chris1221/goldi and we hope that it assists researchers from different domains to better understand and harness the information present in the biomedical literature.

